# Beyond floral visitation: Pollen use reveals selective resource allocation in a generalist solitary bee

**DOI:** 10.64898/2026.05.02.722438

**Authors:** Magda Argueta-Guzmán, Baltazar González, Natalie Van Pelt, Anna C. Dias de Almeida, Trizthan Jiménez Delgado, Lisbeth Peña, Matthew C. Hutchinson, Marilia Palumbo Gaiarsa

## Abstract

A central challenge in characterizing species’ dietary niches is ensuring that foraging data accurately capture the resources used, their relative importance, and how resource abundance shapes foraging patterns. Most studies infer diet breadth and resource-use patterns from observational records, yet such data may lack crucial contextual information documenting how a resource is used or which alternative options were available. For example, in plant-pollinator systems, floral visits involve collecting nectar, pollen, or both, which is allocated to either adult survival or offspring provisioning; visit data cannot distinguish the two, which is important because the resources allocated to offspring ultimately determine reproductive success and population persistence. Here, we experimentally quantified deviation between observed foraging behavior and resources allocated to reproduction by manipulating resource abundance, individually marking bees, and documenting resource use with direct observation, pollen-provision DNA metabarcoding, and pollen-provision microscopy. We show that observed floral visitation patterns did not fully capture the resources allocated to offspring provisioning; pollen from the most frequently visited plants was not the most abundant in provisions. Moreover, we found that resource abundance alone did not consistently explain the observed interaction patterns, challenging the assumption of resource abundance as a strong predictor of plant-pollinator interactions. Our results show that relying on visitation alone may portray a broader niche than is reflected in pollen use, and demonstrate how integrating behavioral foraging observations with offspring requirements can reveal resource-specific patterns that would otherwise remain hidden.

## Introduction

Since Elton’s classic definition of the niche as a species’ functional role, emphasizing trophic interactions (Elton, 1927), ecologists have sought to understand how species use, share, and compete for food resources across heterogeneous environments (Stephens & Krebs, 1986; Bolnick *et al*., 2003; Hengeveld *et al*., 2009; Sponsler *et al*., 2023) and how these processes shape populations and communities (Valdovinos *et al*., 2016). A central challenge in this effort is ensuring that data on foraging behavior accurately capture both the resources used and their relative importance, because biological and methodological biases shape diet records. Resource use can also be measured at different foraging stages, from resource acquisition to consumption and finally as nutrient allocation to growth, survival, or reproduction. Moreover, foraging interactions are inherently dynamic, often difficult to observe, and can shift over seasonal or even weekly timescales as resource abundance changes (Tornberg & Reif, 2006; CaraDonna *et al*., 2017; Daluwatta Galappaththige, 2024). Consequently, controlled settings and a combination of diet-quantification tools are a most likely way to generate accurate data on resource use and to address long-standing questions in trophic ecology, such as the importance of resource availability to diet composition.

In pollinator ecology, estimates of floral resource use are particularly important because they can guide conservation planning and assessments of pollinator species’ vulnerability to environmental change (Burkle *et al*., 2013; Kaiser-Bunbury & Blüthgen, 2015). Floral visitation records are commonly used to characterize pollinator resource use and diet breadth (Fründ *et al*., 2010) but are often unable to differentiate foraging for pollen, nectar, or both, and how resources were subsequently allocated to offspring production. Solitary bees provide a tractable system for investigating whether similar conclusions about resource allocation to reproduction can be drawn from different measures of resource use. Female solitary bees visit flowers to collect nectar, to fuel activity (Cane, 2016; Harano & Sasaki, 2024), and pollen, which they may consume as a protein source (Cane, 2016). However, females also collect pollen and nectar as maternal investment for their offspring, mixing them into pollen-dominated provisions (Danforth *et al*., 2019). Thus, floral visitation cannot be interpreted exclusively as adult feeding or exclusively offspring provisioning and instead, observed floral visits document adults’ foraging interactions with flowers for both purposes, whereas the pollen provisions represent the subset of visited floral resources allocated to offspring production. If the composition of pollen provisions differs substantially from patterns inferred from floral visitation, this could yield contrasting estimates of resource importance, specialization, and diet breadth. Thus, these two approaches may capture related but non-equivalent components of floral resource use. Resolving this question is key to understanding how bees select resources, which has direct applications for conservation planning and assessments of species’ vulnerability to disruptions in plant-pollinator systems (Burkle *et al*., 2013; Kaiser-Bunbury & Blüthgen, 2015).

One potential explanation for why flower visitation and pollen provisions may differ is that they represent different stages of floral resource use. At the visitation stage, adult foraging interactions may be influenced by the relative abundance of blooming plants, as abundance shapes the opportunity for bees to encounter and interact with flowers (Vázquez *et al*., 2007). At the provisioning stage, pollen use may additionally depend on the availability of pollen and the match between bee and floral traits relevant to pollen collection. Whether floral visitation reflects pollen use for offspring provisioning remains largely unexplored. Studies comparing DNA metabarcoding of pollen carried on bees’ bodies with observed floral interactions have generally found broader plant associations from pollen loads (Pornon *et al*., 2017; Encinas-Viso *et al*., 2023; Tourbez *et al*., 2024), with pollen metabarcoding detecting up to twice as many plant species than direct observations of foraging interactions (Arstingstall *et al*., 2021). Although pollen loads represent pollen collected across multiple floral visits during a foraging bout, they do not capture the resources accumulated across repeated foraging trips that are ultimately incorporated into offspring provisions. Comparing observed floral visits with pollen provisions is therefore necessary to determine whether diet breadth based on foraging observations accurately reflects the breadth of resources used to support offspring development.

Comparing flower visitation and pollen provisions depends on the accurate taxonomic identification of pollen grains. Pollen DNA metabarcoding has been widely applied to characterize pollinator floral use across ecological contexts (Milla *et al*., 2022; Casanelles-Abella *et al*., 2022; Arstingstall *et al*., 2023; Ronca *et al*., 2023; Crone *et al*., 2025). Compared with pollen microscopy, metabarcoding can provide greater throughput and finer taxonomic resolution, particularly among morphologically similar pollen grains. However, quantitative estimates from pollen DNA metabarcoding can be biased by variation in marker choice, DNA content per pollen grain, amplification efficiency, and taxon-specific differences in sequence recovery (Bell *et al*., 2019; Swenson & Gemeinholzer, 2021; Moore *et al*., 2022). Independent comparison with microscopy can therefore help assess whether inferred pollen-provision composition reflects biological patterns rather than method-specific bias (Baksay *et al*., 2022), and whether relative abundances estimated from DNA metabarcoding are consistent with pollen-grain counts. Unresolved differences in methodological approaches may otherwise lead to incomplete ecological inferences about pollinator resource use, potentially masking the mechanisms shaping foraging decisions, dietary preferences, and plant-pollinator interactions.

Here, we investigated whether observed patterns of flower visitation by adult female bees reflect the pollen composition of offspring provisions, and how resource abundance shapes visitation patterns. Using experimental plant-bee communities of annual wildflowers and a solitary bee species, we tested whether observed patterns of flower visitation by female bees and pollen provisioned to the offspring yield consistent estimates of plant resource use. Given that pollen varies in its nutritional composition (Roulston *et al*., 2000; Vaudo *et al*., 2024) and potentially in its suitability for larval development (Westreich & Tobin, 2025), we hypothesized that females will selectively incorporate pollen into provisions from only a subset of the plant species they visited while foraging. Using uniquely marked individuals of the solitary bee *Osmia lignaria* in a controlled greenhouse environment, we varied the relative abundance of floral resources across foraging arenas to test (1) whether plant use inferred from observed flower visits was consistent with that reconstructed from pollen provisions. We then explored (2) if variation in resource availability predictably shifted dietary composition. Finally, we evaluated (3) whether two different pollen identification methods—microscopy and DNA metabarcoding—yield quantitatively comparable estimates of pollen-provision composition.

## Materials and Methods

### Study Organism

The blue orchard mason bee (*Osmia lignaria*: Megachilidae) is a generalist, solitary species native to North America and is widely used for pollination services in agricultural systems (Bosch & Kemp, 2001). It has an annual life cycle in which each female constructs a nest consisting of linear series of brood cells, typically in a hollow stem, each containing a single egg and a provision— predominantly made of pollen—to support larval development and metamorphosis. Each cell is sealed with a mud partition before provisioning begins for the next cell. Each provision contains pollen from dozens of foraging bouts (Bosch & Kemp, 2001); collectively, all pollen provisions produced by a female encompass her complete foraging history, as she dies when nesting activity is complete.

### Experimental Design

To investigate whether observed flower visitation patters by adult female bees reflect the pollen composition of offspring provisions of *O. lignaria*, and to determine the relationship between floral abundance and floral resource use, we conducted a controlled greenhouse experiment at the University of California, Merced, in April and May 2025. We first grew from seed four annual wildflower species native to California and commonly visited by *O. lignaria* (Ascher & Pickering, 2020): *Clarkia unguiculata* (Onagraceae), *Nemophila maculata* (Boraginaceae), *Phacelia tanacetifolia* (Boraginaceae), and *Collinsia heterophylla* (Plantaginaceae). We staggered seed sowing to ensure all species would bloom at the same time. Once plants were in full bloom, we established five foraging arenas (80"L x 71"W x 71"H) and placed 40 individually potted plants into each one. All arenas contained all four focal plant species, but arenas differed in each species’ relative abundance (Fig. 2a; Table S1). We kept the temperature at 27*^◦^*C. Into each foraging arena, we released 12 males and five individually marked females, reflecting the sex ratio observed in wild populations and commonly used for captive rearing in this species (Bosch & Kemp, 2001). We placed one nest block in each arena containing multiple individual paper straws (simulating hollow stems) and supplied moist clay into each arena to facilitate nesting (Bosch & Kemp, 2001).

### Observed Floral Visits and Female Tracking

After a two-day acclimation period, we monitored the flower-visitation patterns of the five females in each arena daily for 20 days (Fig. 1). Each day, we recorded each female’s flower visits during one 15-minute focal observation period. To account for diel variation in foraging patterns, we randomized observation times between 10 am and 3 pm when bees are most active. We defined a visit as a foraging interaction when we observed a female touching the flower’s anthers and/or probing the base of the corolla. For each visit, we documented the female’s identity and the plant species. Finally, for each female in each foraging arena, we quantified floral-visitation patterns as the total number of visits to each plant species across the observation period.

**Figure 1:**
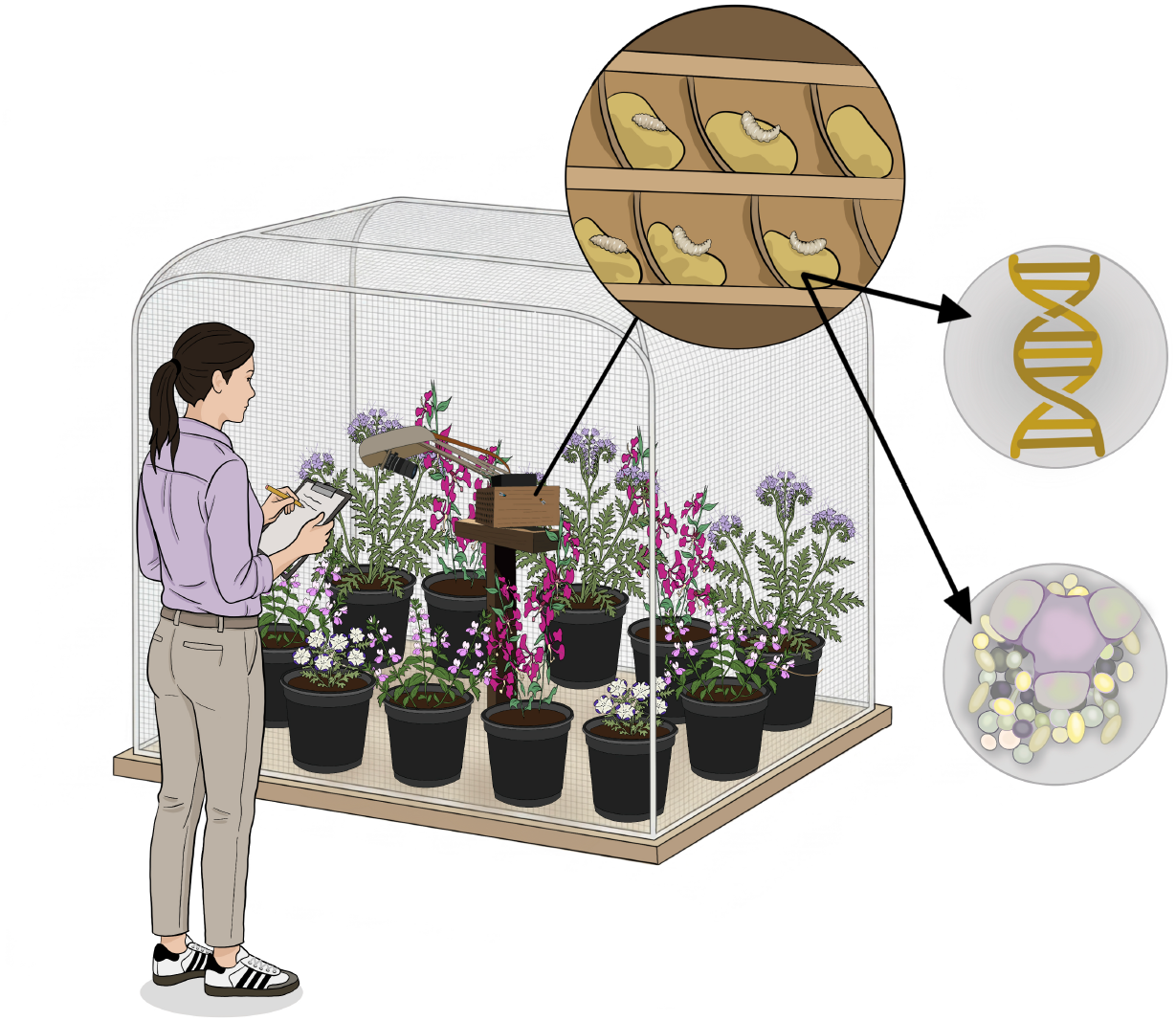
Experimental set-up used in this study. We established five foraging arenas (80"L x 71"W x 71"H) containing 40 plants of four California native annual plant species commonly visited by the solitary bee *Osmia lignaria*: *Clarkia unguiculata, Nemophila maculata, Phacelia tanacetifolia* and *Collinsia heterophylla*. Each arena included a nest block and moist clay to facilitate nesting. We monitored daily bee–flower interactions for 20 days to quantify floral visitation patterns. We marked individual females to link foraging patterns to nesting, and continuously video-recorded the nest entrance to assign nests to specific females. We collected completed nests to characterize pollen provisions using both DNA metabarcoding (ITS2 rRNA region) and pollen microscopy. Illustration by NvP.

In addition, we continuously video-recorded nest entrances throughout the experiment to assign nesting straws to specific females (Fig. 1). We inspected each nest block daily for completed nest straws (i.e., those with a sealed entrance) and immediately transferred completed provisions (n = 55) to a -80°C freezer for storage and DNA preservation before molecular and microscopic analyses.

### Pollen Metabarcoding

We performed DNA metabarcoding on all obtained pollen provisions (n = 55) in triplicate, using the universal ITS2 primers (White *et al*., 1990; Chen *et al*., 2010), to assess the pollen composition of each pollen provision. We selected ITS2 after *in silico* comparisons showed that it distinguished all focal plants to at least the genus level (i.e., sufficient to distinguish all plants used in the experiment), whereas *rbcL* could not differentiate two genera and assigned both only to their shared family. To extract DNA from each provision, we carefully dissected each straw by making a longitudinal cut with clean utensils under sterile conditions and transferred 50 mg of each pollen provision (a pea-sized subsample) into sterile vials labeled with the corresponding foraging arena, straw, and pollen provision number. We extracted DNA from each pollen provision using the Qiagen Plant Pro kit (Qiagen, MD, USA) and following the manufacturer’s protocol after first disrupting and homogenizing pollen grains by sonicating them with 0.5 mm zirconia beads at 30 Hz for 4 minutes in a TissueLyser LT. We included reagent-only blanks as negative controls and DNA from *Epipremnum* spp. (Araceae) as positive controls. DNA concentrations ranged from 40 to *>* 100 ng *µ*L*^−^*^1^, as measured using the Qubit dsDNA High-Sensitivity Assay (Thermo Fisher Scientific, Waltham, MA, USA). To prepare libraries for pollen DNA metabarcoding, we used a dual-index inline barcoding approach following Kembel *et al*. (2014), in which primers were composed of an Illumina sequencing primer (forward or reverse), an eight-nucleotide barcode, and the forward or reverse genomic oligonucleotide targeting the internal transcribed spacer 2 (ITS2 rRNA) plant marker. We amplified 329–374 bp using the ITS2-S2F (ATG CGA TAC TTG GTG TGA AT) (Chen *et al*., 2010) and ITS2-S4R (TCC TCC GCT TAT TGA TAT GC) (White *et al*., 1990) primer pair. For each reaction, we used 10 µl ultrapure water, 10 µl Phusion MasterMix (Thermo Scientific, Baltics UAB), 1 µl of each primer (10 µM), and 4 µl of DNA. We used a 58°C annealing temperature, 35 cycles, and included negative PCR controls. We cleaned the first PCR reaction enzymatically using ExoSAP-IT Fast (Thermo Scientific) according to the manufacturer’s protocol. We then used 1 µl of the clean PCR product as a template for a second PCR, using HPLC-purified primers to complete the Illumina sequencing construct as in Kembel *et al*. (2014): CAA GCA GAA GAC GGC ATA CGA GAT CGG TCT CGG CAT TCC TGC and AAT GAT ACG GCG ACC ACC GAG ATC TAC ACT CTT TCC CTA CAC GAC G. For these reactions, we used a 58°C annealing temperature, 15 cycles, and negative and positive controls from the first PCR. All positive controls yielded amplicons of the expected size for both PCR steps. To standardize DNA content across libraries, we normalized 18 µl of the second PCR product using SequalPrep normalization plates (Invitrogen, MA) following the manufacturer’s protocol. We established a library pool by combining 5 µl from each normalized library. To eliminate primer dimers and excess master mix components, we conducted library purification using AMPure XP beads (Beckman Coulter, CA). We also tested within-sample consistency by preparing and amplifying libraries with different index pairs in triplicate for each provision. We evaluated our library fragment size distributions on a Fragment Analyzer (Advanced Analytical Technologies, NY) before sequencing on an Illumina MiSeq i100 series instrument, using the V3 2 × 300 reagent kit, a process carried out by the Vincent J. Coates Genomics Sequencing Lab at the University of California, Berkeley.

We then performed bioinformatics and taxonomic assignment by demultiplexing raw reads in QIIME 2 (version 2024.10) (Bolyen *et al*., 2019) and trimming adapters and primers with the Cutadapt plugin using default settings (Martin, 2011). We assessed read quality with FastQC (version 0.12.1) (Andrews, 2010) and summarized the results with MultiQC (version 1.28) (Ewels *et al*., 2016). To denoise and merge reads, remove chimeras, and generate amplicon sequence variants (ASVs), we used the DADA2 pipeline through q2-dada2 (Callahan *et al*., 2016). We allowed a maximum of two and three expected errors in forward and reverse reads, respectively, and used consensus chimera removal with a minimum parent-over-abundance ratio of 2 (Callahan *et al*., 2016). Across the dataset, DADA2 processed 14,426,923 input reads. Quality filtering retained 94.4% of reads, 87.4% of input reads were successfully merged, and 10,258,534 non-chimeric reads remained (71.1% of input reads and 81.4% of merged reads; Table S2, S3, Fig. S1, and Fig. S2). To standardize sequencing effort across samples, we rarefied our sequenced libraries by random subsampling to 26,133 reads, where our accumulation curves flatten while retaining all samples and all but two technical triplicates. We exported this rarefied feature table for further ecological analyses and averaged triplicate composition for each provision and individual female. We retrieved ITS2 sequences from NCBI (November 04, 2025) for the genera of interest (*Nemophila, Clarkia, Collinsia*, and *Phacelia*) and created a BLAST reference database with makeblastdb (Camacho, 2024). To assign taxonomy to the obtained ASVs, we conducted local BLAST searches (Altschul *et al*., 1990; Camacho *et al*., 2009) against our custom curated reference database. We used a minimum genus percent identity of 0.95 and saved the top matches with *e*-values <1e-20, selecting the match with the highest bit score. Using these criteria, we assigned 98.6% of total rarefied reads to genus level (Table S4). Because each focal genus was represented by a single species in our dataset, our taxonomic assignments also identified corresponding plant species.

### Pollen Microscopy

To test whether pollen composition and relative proportions inferred from DNA metabarcoding reflected those from pollen-grain quantification under a microscope, we prepared reference slides for the four focal plant taxa at 400x and 600x magnification using sunflower oil as the mounting medium. For each pollen provision, we first thoroughly homogenized the sample and then mounted a subsample (5 µl) on a slide. We overlaid each field of view with a grid divided into four equal-area quadrants. Four independent observers identified and counted pollen grains at 400x magnification in two randomly selected quadrants per slide, with a mean of 70 grains per slide. To assess interobserver consistency, we quantified variation among observers. For subsequent analyses, we used mean counts across observers for each sample and treated these values as the relative abundance of each plant species in each pollen provision.

### Statistical Analyses

To evaluate whether resource use differed between observational and provision-based approaches, we analyzed compositional variation among the three methods (observation, metabarcoding, and microscopy) using a permutational multivariate analysis of variance (PERMANOVA; Anderson (2001)), implemented with the adonis2 function in *vegan* (Oksanen *et al*., 2025). We constrained permutations within females (stratified by female identity) to account for repeated measures. To verify that significant PERMANOVA results were not driven by differences in multivariate dispersion among methods, we tested for homogeneity of dispersion using the function betadisper from *vegan* (Oksanen *et al*., 2025). We used a non-metric multidimensional scaling (NMDS) ordination to visualize multivariate diet structure across methods. Furthermore, for each female within each foraging arena, we quantified dissimilarity between these measures as Bray-Curtis distance (Bray & Curtis, 1957), calculated on relative abundances with the function vegdist from the *vegan* package (Oksanen *et al*., 2025), which provided a per-individual dissimilarity measure ranging from 0 (identical composition and proportion) to 1 (no shared floral taxa in the diet).

To assess whether visitation- and pollen-based resource estimates reflected floral availability, we calculated the Bray-Curtis dissimilarity between each female’s diet composition and the floral composition of her foraging arena. We estimated each female’s diet composition from observed flower visits and pollen metabarcoding (n = 16), and used plant relative abundances as a measure of arena-level floral availability. We then modeled Bray-Curtis dissimilarity as a function of diet quantification method using a generalized linear mixed-effects model in glmmTMB (Brooks *et al*., 2017), with a beta error distribution and logit link. We included quantification method as a fixed effect and tested alternative random-intercept structures for arena, female identity, or both. We compared candidate models using the Akaike Information Criterion (AIC) and singularity tests with a tolerance of 1e-05 (Fox *et al*., 2015). The best model, supported by the AIC, was *Y ∼ method* + (1*|arena*), including the random intercept for the arena (also not singular). We tested scaled residual dispersion and outliers using DHARMa (Hartig, 2026) and found no evidence of over- or under-dispersion (*p* = 0.54), outliers (*p* = 1.0), or deviation from uniformity (*p* = 0.31).

Moreover, within each foraging arena, we also quantified deviation from expected resource use for each female’s observed flower visits and for each provision. We used one-sample t-tests to evaluate whether the mean deviation of diet from resource availability was significantly greater than zero. No deviation from zero indicates that bees track the floral availability of that resource, whereas positive or negative values indicate over- or under-representation of a specific flower species relative to its availability.

To test whether pollen identification via DNA metabarcoding and microscopy yields comparable quantitative estimates of pollen provision composition (n = 55), we assessed the correlation between the relative pollen abundance of each plant species in each provision using Spearman rank correlation. We also evaluated compositional consistency in DNA metabarcoding results across triplicates to assess potential stochasticity in library preparation and amplification. After calculating pairwise Pearson correlations among triplicates for each pollen provision and finding high internal consistency (minimum mean Pearson correlation across replication pairs *r* = 0.973, *SD* = 0.034, Fig. S3), we averaged triplicates by sample for subsequent analyses. We conducted all analyses in R version 4.5.2 (R Core Team, 2025) and all figures were prepared with ggplot2 from tidyverse (Wickham *et al*., 2019) and patchwork (Pedersen, 2025).

## Results

We observed 545 interactions between female bees and the four plant species across 1,500 minutes of observations. Across all arenas, *P. tanacetifolia* received the most visits (n = 198; 36.3%), followed by *C. unguiculata* (n = 143; 26.2%), and *C. heterophyla* (n = 140; 25.7%). *N. maculata* received the fewest visits (n = 64; 11.7%). We recorded the most visits in the *N. maculata*-dominated arena (n = 187), followed by the *C. heterophylla*-dominated arena (n = 178), and the fewest in the *P. tanacetifolia*-dominated arena (n = 51). Of the 25 females in the experiment, 32% nested (n = 8), yielding a total of 55 pollen provisions across our experimental foraging arenas (provisions per female, mean = 6.9, range = 1-12). At the end of the experiment, we found 51 provisions (92.7%) containing offspring, including 20 eggs and 31 viable larvae, and four containing no offspring (Table S5).

We found that diet composition inferred from observed flower visits (Fig. 2b) differed substantially from that reconstructed from pollen provisions, regardless of the pollen-identification method (Fig. 2c-d). Diet-quantification method was a significant predictor of dissimilarity in diet composition (*F* = 6.2, R^2^ = 0.37, *p* = 0.001; Figs. 3a-c), which was not driven by differences in within-group dispersion (betadisper: *F* = 2.11, *p* = 0.14). These results suggest that diets inferred from each method differ in composition rather than in variance. For instance, *C. unguiculata* appeared in diets far more frequently when diet was quantified from observed visits than in either pollen-based method (Fig. 2b-d), whereas *C. heterophylla* was detected at higher relative abundance in pollen provisions than in visitation data. Per-female Bray-Curtis distances differed among method-pair categories; both pollen-based methods showed high dissimilarity from observed floral visitation patterns (mean = 0.37-0.46; Figs. 3d-e), but were similar to one another (mean = 0.18; Fig. 3f).

**Figure 2:**
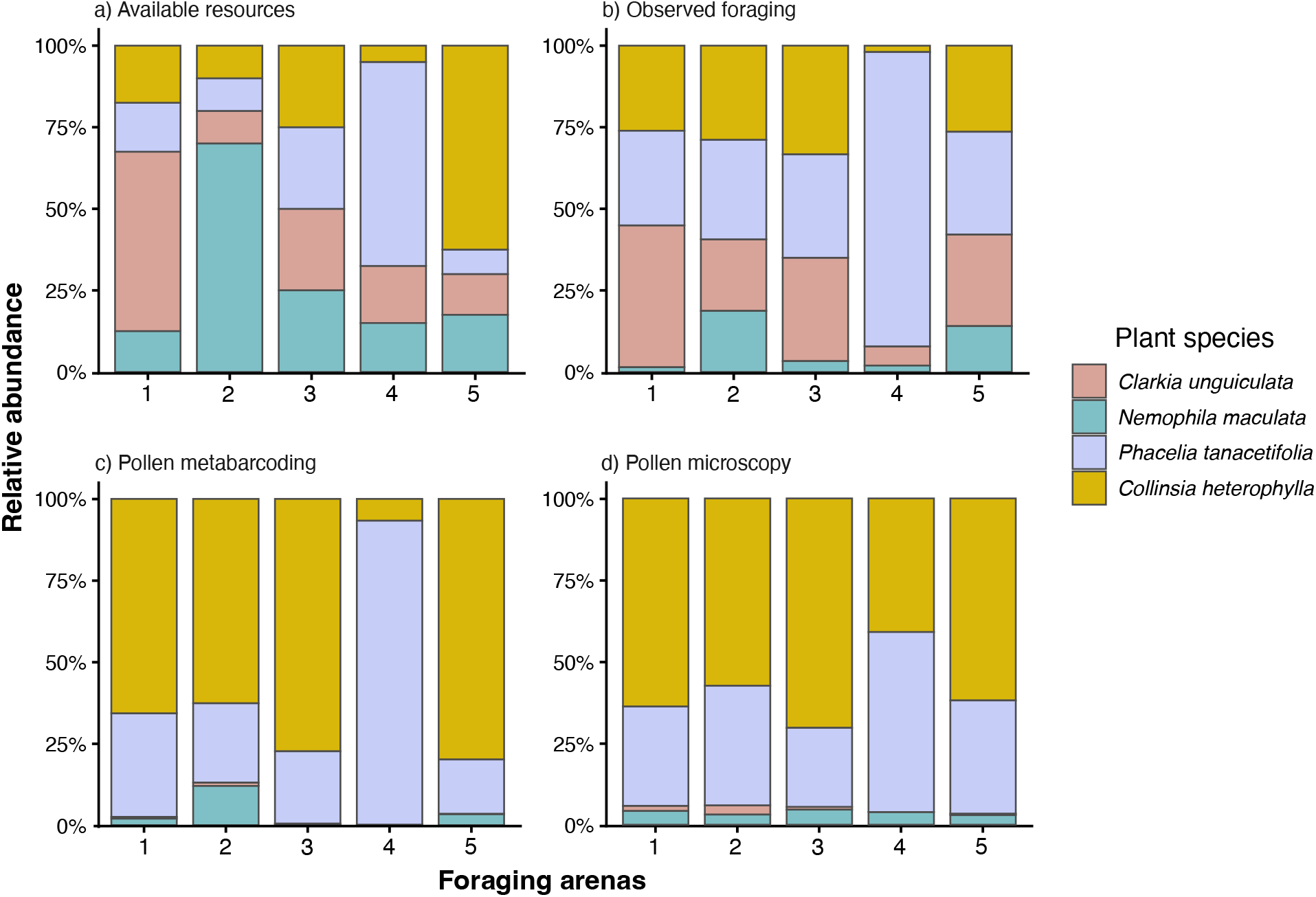
Relative abundance of floral resources and pollen use by *Osmia lignaria*. Within each arena, we calculated percentages as follows: (a) the number of flowering plants of each focal species divided by the total number of flowering plants; (b) visits to each plant species divided by the total number of observed flower visits; (c) the proportion of assigned ITS2 reads for each plant species per provision; and (d) the proportion of pollen grains for each plant species per provision. We combined provision-level proportions within each arena and rescaled them to sum to one. *N. maculata* was detected by metabarcoding in all arenas (Table S6), although its low relative abundance in arena 3 and 4 makes its presence difficult to discern visually in panel (c).

**Figure 3:**
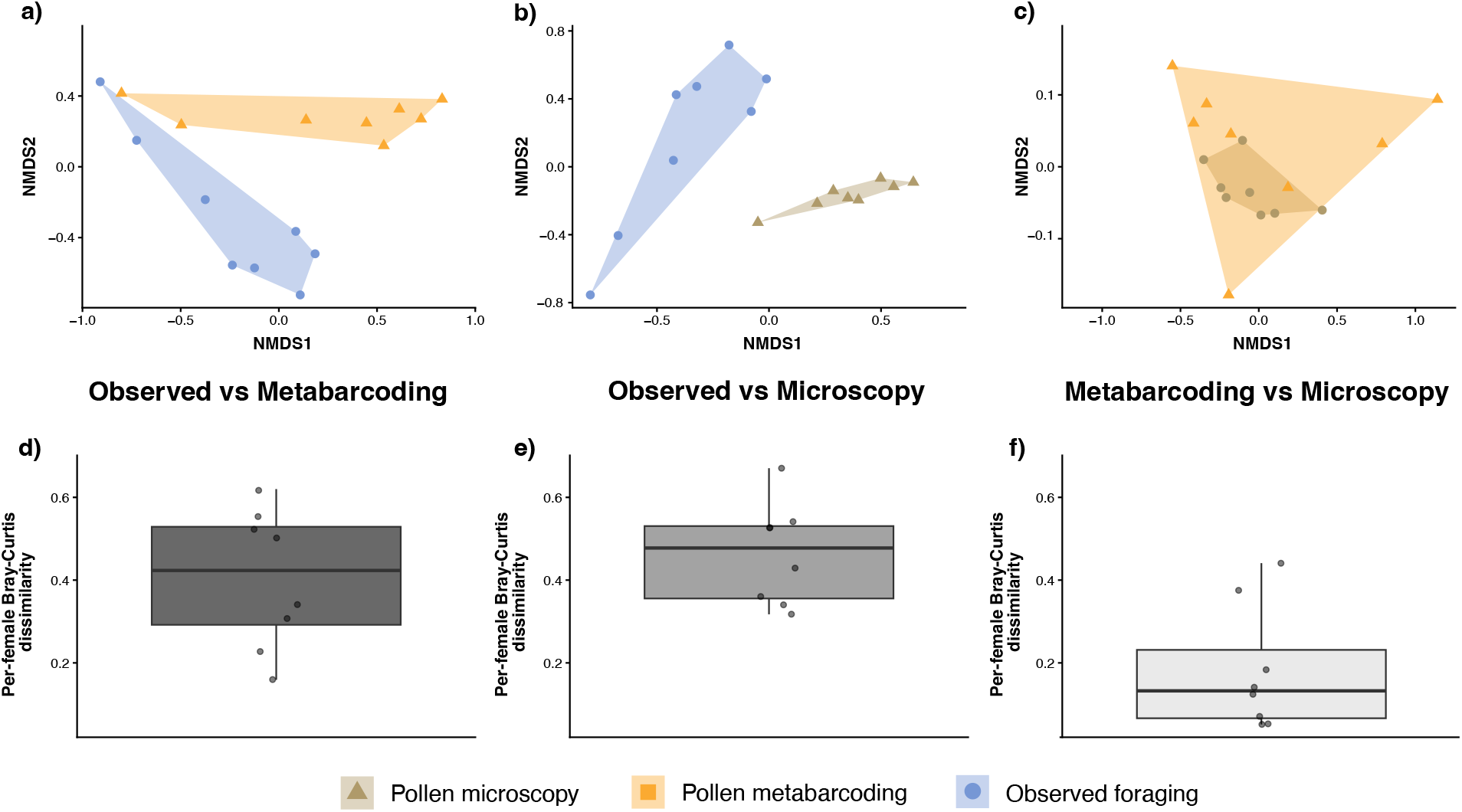
Correspondence among methods used to reconstruct *Osmia lignaria* diets. Panels (a–c) show non-metric multidimensional scaling (NMDS) ordinations (n = 8 females), and panels (d–f) show per-female Bray–Curtis dissimilarities between each method pair. Comparisons include observed floral visits vs. pollen DNA metabarcoding (a, d), observed floral visits vs. pollen microscopy (b, e), and pollen DNA metabarcoding vs. pollen microscopy (c, f).

Resource use estimates based on observed flower visits—reflecting nectar and pollen foraging— were more compositionally similar to resource availability than those based on DNA metabarcoding— capturing only pollen incorporated into provisions (Bray-Curtis mean = 0.316 and 0.5, respectively; Fig. 4). Our generalized linear mixed-effects model supported this stronger effect of resource availability on diets reconstructed from observed flower visits than diets reconstructed from pollen provisioning (*β* = 0.76 *±* 0.30 SE, *p* = 0.01). However, floral availability alone still did not fully explain observed flower visits. Females showed consistent deviations from nonselective for-aging. For example, observed foraging activity on *P. tanacetifolia* was more frequent than expected based on its availability (mean difference = 0.17, *p <<* 0.01; Fig. 2a and 2b), whereas *N. maculata* was visited less frequently than expected (mean difference = *−*0.22, *p <<* 0.01; Fig. 2a and 2b). In contrast, visitation to *C. unguiculata* and *C. heterophylla* were similar to expectations based on resource availability (mean difference = 0.04, *p <<* 0.01 and mean difference = 0.009, *p <<* 0.01, respectively; Fig. 2a and 2b), indicating that the bees’ foraging effort, at least for nectar, tracked the abundance of these plants. These results suggest that foraging females preferentially visited *P. tanacetifolia*, but visited *C. unguiculata* and *C. heterophylla* in proportion to their availability.

**Figure 4:**
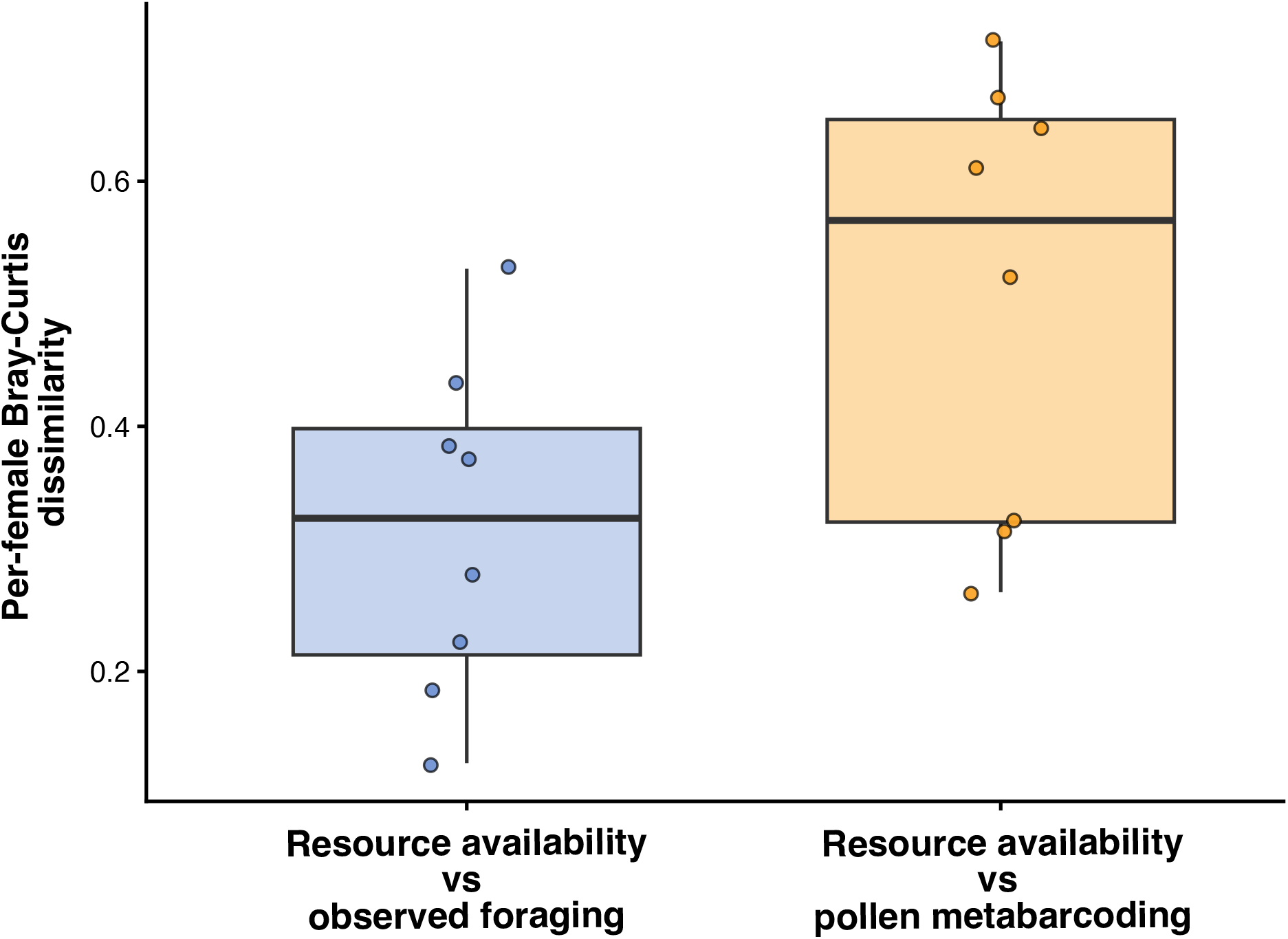
Diet estimates differ in their correspondence with floral resource availability across methods used to characterize *Osmia lignaria* diets (Bray-Curtis dissimilarity per female). Observed floral visits capture both nectar and pollen foraging, whereas pollen metabarcoding reflects only pollen used for offspring provisioning. This pattern was supported by a generalized linear mixed model with beta error distribution and logit link (*β* = 0.76 *±* 0.30 SE, *p* = 0.01).

For pollen provisions, we found *P. tanacetifolia* and *C. heterophylla* were overrepresented in provisions relative to their availability (mean difference = 0.16, *p <<* 0.01 and mean difference = 0.33, *p <<* 0.01, respectively; Fig. 2a and 2c). In contrast, *C. unguiculata* was substantially under-represented (mean difference = -0.30, *p <<* 0.01), as was *N. maculata* (mean difference = -0.20, *p <<* 0.01; Fig. 2a and 2c). These patterns indicate selectivity in pollen foraging, where pollen provisions were not assembled in proportion to floral availability, with *P. tanacetifolia* and *C. heterophylla* contributing disproportionately to offspring provisions, regardless of their availability. Microscopy estimates were generally consistent among observers, with mean absolute differences in relative abundance ranging from 0.03 to 0.13 across the four plant taxa. We found strong correlations between DNA metabarcoding (ITS2) and microscopy estimates of relative pollen abundances in provisions for *C. heterophylla* (*ρ* = 0.77), *P. tanacetifolia* (*ρ* = 0.72), and *C. unguiculata* (*ρ* = 0.64), but the two were negatively and weakly correlated for *N. maculata* (*ρ* = -0.2; Bonferroni-adjusted *p <* 0.01 in all cases; Figs. 2c-d and Table S6).

## Discussion

Here, we experimentally manipulated resource abundance in a greenhouse system to evaluate how resource use varies between observed floral visitation and resources ultimately allocated to offspring using the solitary bee *O. lignaria*. Integrating three different approaches to quantify floral resource use, we identified clear differences between visitation-based and provision-based estimates. By examining how resource availability influences adult foraging, we found that plant abundance affects flower visitation patterns more strongly than pollen provisioning patterns. The greater selectivity observed in pollen provisioning suggests that factors other than availability (*e.g.,* nutritional suitability) may influence which pollen resources are incorporated into offspring provisions. Yet, frequently visited plants that contribute little pollen to provisions may still provide nectar for adult self-maintenance and for mixing the pollen provisions. In addition, resource availability alone was insufficient to fully predict diet composition from either method, suggesting other factors also shape foraging decisions. Our results further revealed that diets inferred from adult floral-visitation patterns and those derived from pollen provisions differed substantially. Finally, we found that two pollen analysis methods, DNA metabarcoding and microscopy, yielded broadly consistent estimates of bee diets. Overall, our results highlight that integrating observational data with pollen-provision analyses may provide a more complete picture of floral resource use. Because offspring provisioning directly determines offspring survival and recruitment, relying solely on floral visitation data may underestimate the importance of specific resources for bee populations. We show that integrating both measures of resource use (adult foraging and pollen provisioning) can advance our understanding on the importance of specific resources for bee populations.

We found that visitation-based estimates included floral resources that were nearly absent from offspring provisions, whereas provision-based estimates highlighted a narrower subset of resources allocated to offspring. Adult flower visitation patterns do not necessarily overestimate the complete dietary niche of *O. lignaria*, but our results suggest that they do reveal that the range of plants visited can be broader than the range contributing substantially to provisioning pollen. One consequence of this mismatch in diet estimates is that flower visits and pollen provisions can provide complementary descriptions of floral resource use: some visited plants may primarily contribute resources for adult maintenance, whereas others can be relatively more important for offspring provisioning. Therefore, bee species classified as generalists based on visitation data may still exhibit functional specialization (Bolnick *et al*., 2003) if they use different subsets of floral resources for distinct foraging goals. As a result, apparent specialization in species classified as generalists may emerge from different drivers of foraging decisions throughout their lifetimes.

One plausible mechanism underlying this apparent diet breadth mismatch is resource-specific foraging: females may obtain nectar from a broad range of plants while preferentially collecting pollen from species that may exhibit either more nutritionally suitable pollen or are easier to harvest. For example, *C. heterophylla* and *P. tanacetifolia* were both frequently visited and present in provisions, whereas *C. unguiculata* received numerous visits but was nearly absent in pollen provisions. The high relative abundance of *C. heterophylla* and *P. tanacetifolia* in provisions confirms their use as pollen sources but does not rule out nectar collection during visits to these flowers. Thus, although we did not measure nectar consumption directly, floral visits to *C. unguiculata* flowers may primarily reflect nectar foraging, which could explain why *C. unguiculata* dominates visitation records yet is almost absent from pollen provisions. Moreover, floral abundance may not reflect collectible pollen, and the viscin-rich, strongly aggregated pollen of *C. unguiculata* may be difficult for *O. lignaria* to collect and pack (Nemeth & Smith-Huerta, 1997). Consistent with a potential role as a nectar source, *C. unguiculata* can produce substantially more nectar and sucrose than *P. tanacetifolia*, depending on watering conditions (Gregor, 2025). Thus, differences in reward function, handling requirements, and trait complementarity may explain why females visited a broader range of plants than those from which they collected pollen. Recent studies have also found incomplete correspondence between pollen loads in bees’ bodies and either foraging observations (Pornon *et al*., 2017; Encinas-Viso *et al*., 2023) or pollen ultimately incorporated into pollen provisions (Klečka *et al*., 2022; Crone *et al*., 2023). Such selective pollen use may reduce pollen-transport efficiency for particular plant species when otherwise generalist bees concentrate pollen in pollen-carrying structures, making it less available for transfer to conspecific stigmas (Weinman *et al*., 2023). Whether this translates into a decrease in plant reproductive success remains to be tested. Taken together, our results suggest that pollen analyses alone may overlook frequently visited plants that contribute little pollen but provide nectar for self-maintenance and/or offspring provisioning.

When estimating resource use based on observed floral visits rather than on pollen provisions, we found that observed floral visits were more closely aligned to resource availability, indicating that although adult females visit a broad set of plants, pollen allocation into nest provisions is more selective. Together with flower abundance, phenology is another important factor shaping plant-pollinator interaction patterns(Vázquez *et al*., 2007; Labonté *et al*., 2023). Here, we controlled for floral abundance and removed the effect of phenology by ensuring all plants bloomed simultaneously throughout the study. We found that less abundant plant species could contribute disproportionately to pollen provisions, indicating that processes beyond abundance may play an important role in shaping plant-pollinator interactions, particularly patterns of pollen use. Selective provisioning has been documented in other generalist *Osmia* and bumblebee species (Sedivy *et al*., 2011; Haider *et al*., 2013; Vanderplanck *et al*., 2018), as well as avoidance of pollen that can be detrimental to larval development (Moerman *et al*., 2017). Thus, larval nutritional requirements may influence which plant species are represented in the pollen used for offspring provisioning.

Overall, we found that pollen metabarcoding and microscopy methods produced similar compositional data for *P. tanacetifolia*, *C. heterophylla*, and *C. unguiculata*. Although pollen metabar-coding detected *N. maculata* in all five arenas, we found a weaker correlation with microscopy methods for this plant species, which was generally at low relative abundance Table S6. Although ITS2 metabarcoding can reliably reconstructs pollen community composition (Sickel *et al*., 2015), variation in ribosomal DNA copy number can affect quantitative estimates (Lofgren *et al*., 2019; Bell *et al*., 2019). Our choice to use metabarcoding for our ecological analyses was supported by the performance of our curated ITS2 reference database, which distinguished all four focal species, assigned 98.6% of rarefied reads, and yielded highly consistent results across PCR triplicates. The high technical repeatability of metabarcoding therefore provided a standardized basis for comparing pollen provisions. Our results suggest that relying on a single pollen-provision sample is sufficient to accurately characterize pollen composition in experimental settings. However, when using DNA metabarcoding to characterize pollen provisions when the available species pool is unknown, PCR replicates and local reference DNA libraries will still be advantageous given their increased likelihood of detecting rare taxa and achieving acceptable taxonomic resolution (Shirazi *et al*., 2021).

## Conclusion

Our results show that floral resource use encompasses complementary dimensions of a pollinator’s niche. We found that observed flower visits differed from pollen in offspring provisions, highlighting the importance of integrating different stages of floral resource use when characterizing pollinators’ dietary breadth, as methodological choices can shape our understanding of pollinators’ use of floral resources. Moreover, we found that resource availability alone did not fully explain either observed floral visits or the composition of resources allocated to nests. However, observed floral visitation more closely tracked local floral availability, whereas pollen provisioning showed greater selectivity. While microscopy and DNA metabarcoding methods generated similar estimates of pollen-provision composition, integrating pollen-based approaches with direct foraging observations revealed otherwise hidden dimensions of floral resource use. Thus, plant-pollinator interaction networks constructed from flower visits may not fully represent the relative importance of plants as pollen sources, potentially overlooking important resources that sustain offspring development and, in turn, population persistence. Building on nutritional landscape approaches (Stephen & Rehan, 2026; Vaudo *et al*., 2024), future studies should determine how pollen and nectar quality influence foraging, provisioning, and reproductive success. Connecting these components will provide a more complete understanding of how floral communities support pollinator populations.

## Acknowledgments

We thank Paulo Roberto Guimarães Jr. and members of the Guimarães Lab for valuable feedback on earlier versions of this manuscript. We acknowledge the use of artificial intelligence tools to improve grammar and language clarity, but not to generate text, scientific content, or analysis, or to assist with data interpretation. We thank two anonymous reviewers for their helpful feedback.

## Funding

MAG, BG, and MPG thank the support of the California Natural Resources Agency, and MPG acknowledges the support of the National Science Foundation under Grant No. 2437073.

## Data Availability Statement

DNA metabarcoding sequences (ITS2) from *Phacelia tanacetifolia* (Boraginaceae), *Nemophila maculata* (Boraginaceae), *Collinsia heterophylla* (Plantaginaceae), and *Clarkia unguiculata* (Onagraceae) have been uploaded to the NCBI nucleotide database (SUB16135792).

## Author Contributions

MAG, MPG, and MCH conceived the idea. MPG designed the empirical experiment, and MAG, BG, NVP, ACDA, TJD, and LP implemented the experiment and collected foraging observation data. MAG conducted pollen DNA lab work and analyses and led the pollen identification through microscopy, assisted by NVP, ACDA, TJD, and LP. MAG analyzed results, prepared the figures, and wrote the first draft of the paper, with input from MPG and MCH. MPG, MCH, and BG gave feedback on the final draft, and all authors gave final approval for submission.

## Supporting information

### Floral composition of our experimental arenas

**Table S1:** Floral composition of experimental foraging arenas. Values indicate the number of flowering individuals per species in each treatment (total = 40 plants per arena) for the plants *Clarkia unguiculata* (Onagraceae), *Nemophila maculata* (Boraginaceae), *Phacelia tanacetifolia* (Boraginaceae), and *Collinsia heterophylla* (Plantaginaceae).

| Arena | *C. unguiculata* | *P. tanacetifolia* | *C. heterophylla* | *N. maculata* | Total |
| --- | --- | --- | --- | --- | --- |
| 1 | 22 | 6 | 7 | 5 | 40 |
| 2 | 4 | 4 | 4 | 28 | 40 |
| 3 | 10 | 10 | 10 | 10 | 40 |
| 4 | 7 | 25 | 2 | 6 | 40 |
| 5 | 5 | 3 | 25 | 7 | 40 |

### Bioinformatics validation

#### Sequencing processing, quality filtering, and chimera detection details

We processed paired-end ITS2 reads with DADA2 through q2-dada2 in QIIME 2 (version 2024.10) (Callahan *et al*., 2016). We removed no additional bases from the 5*^′^* ends after primer trimming, truncated forward and reverse reads at 270 and 260 bp, respectively, and allowed maximum expected-error values of two for forward reads and three for reverse reads. We used a truncation quality score of two and the consensus method for chimera removal.

We evaluated sensitivity to the minimum fold-parent-over-abundance ratio used for chimera identification. The default analysis used a ratio of one, whereas the sensitivity analysis used a ratio of two; all other DADA2 settings and input reads remained unchanged. Quality filtering retained 94.4% of 14,426,923 input reads, and 12,604,894 reads merged successfully (87.4% of input reads). The ratio-one setting retained 8,846,900 non-chimeric reads, whereas the ratio-two setting retained 10,258,534 reads (Table S2). Thus, the ratio-two setting retained 1,411,634 additional reads while preserving the same preceding filtering, denoising, and merging outcomes.

At the sample level the ratio-two setting increased read retention consistently across the sequenced libraries (Fig. S1). Every library retaining less than 60% of merged reads under the ratio-one setting retained more than 60% under the ratio-two setting. We interpreted this pattern together with the independent sequence checks described below rather than treating read retention alone as evidence that individual ASVs were true biological sequences.

**Table S2:** Read retention and feature recovery for all the sequenced libraries under the ratio-one and ratio-two DADA2 chimera settings for minimum fold-parent-over-abundance. Percentages are pooled across libraries.

| Metric | Ratio = 1 | Ratio = 2 |
| --- | --- | --- |
| Input reads | 14,426,923 | 14,426,923 |
| Reads passing quality filtering (%) | 94.4 | **94.4** |
| Merged reads | 12,604,894 | 12,604,894 |
| Merged reads/input reads (%) | 87.4 | **87.4** |
| Non-chimeric reads | 8,846,900 | 10,258,534 |
| Non-chimeric reads/merged reads (%) | 70.2 | **81.4** |
| Non-chimeric reads/input reads (%) | 61.3 | **71.1** |
| Number of ASVs | 731 | 967 |
| Libraries with at least 26,133 reads | 168 | 175 |

#### Audit of ASVs chimeras

As part of our quality control pipeline, we audited the ASVs reported as chimeras from the two-settings ratio described above. The ratio-two analysis recovered 236 GH ASVs that were absent under the ration-one chimera setting. We evaluated the most abundant restored ASVs using full-length local BLAST matches, their distribution among samples and technical replicates, and an independent *de novo* VSEARCH chimera screen. These inspections indicated that the one-ratio setting threshold removed at least some well-supported biological sequences of *Phacelia tanacetifolia* and *Collinsia heterophylla*. **Thus, we used the ratio-two setting for the primary DADA2 analysis and subsequent ecological analyses. These sensitivity analyses indicated that erroneous classification of valid ASVs as chimeric was unlikely to have affected our conclusions.**

Despite recovering 236 ASVs, the ratio-two setting produced only minor changes in the relative abundances of the focal plants in comparison to the ratio-one setting. The mean absolute difference between settings was 1.93 percentage points at both the provision and arena levels, while the maximum absolute differences were 9.21 and 5.47 percentage points, respectively (Table S3). The dominant compositional patterns and downstream ecological conclusions were unchanged regardless of using chimera ratio-one or ratio-two setting. **The dominant compositional patterns and downstream ecological conclusions were consistent across both DADA2 chimera-detection settings.**

#### Taxonomic assignment and rarefaction

We assigned taxonomy by conducting local BLAST searches against a custom reference database containing ITS2 sequences for the four focal species (Each belonging to a different genus). We retained genus-level matches with at least 95% sequence identity and *e*-values *<* 1 *×* 10*^−^*^20^ and selected the qualifying match with the highest bit score. Regarding negative controls, they had very low number of reads and didn’t pass the quality filters.

**Table S3:** Sensitivity of community composition to the DADA2 parent-over-abundance ratio.

| Metric | Value |
| --- | --- |
| ASVs under the ratio-one setting | 731 |
| ASVs under the ratio-two setting | 967 |
| ASVs restored under the ratio-two setting | 236 |
| Reads restored under the ratio-two setting | 1,411,634 |
| Restored reads as a percentage of ratio-two reads | 13.8 |
| Mean absolute provision-level change (percentage points) | 1.93 |
| Maximum absolute provision-level change (percentage points) | 9.21 |
| Mean absolute arena-level change (percentage points) | 1.93 |
| Maximum absolute arena-level change (percentage points) | 5.47 |
| Reads from the restored *Nemophila* ASV | 28,969 (2%) |
| Percentage reads from the restored belonging to *N. maculata* | 2% |
| Percentage reads from the restored belonging to *P. tanacetifolia* and *C. heterophylla* | 98% |

To standardize sequencing effort at the sample level, we rarefied each library to 26,133 reads per library without replacement, using a fixed random seed of one. This procedure produced 4,573,275 reads before taxonomy-based filtering. Of these, 4,508,488 reads (98.6%) met the genus-level assignment criterion and were retained for ecological analyses; 64,787 reads (1.4%) remained unassigned under this criterion (Table S4).

**Table S4:** Final dataset after DADA2 pipeline.

| Metric | Value |
| --- | --- |
| GH libraries before rarefaction | 165 |
| GH libraries retained | 165 |
| Rarefaction depth per library | 26,133 |
| Total reads after rarefaction | 4,573,275 |
| Reads assigned to genus | 4,508,488 |
| Reads assigned to genus (%) | 98.6 |
| Reads not meeting the assignment criterion | 64,787 |
| Reads not meeting the assignment criterion (%) | 1.4 |

**Table S5:** Reproductive output of *Osmia lignaria* in experimental foraging arenas. Numbers and percentages of eggs, viable larvae, and provisions containing no offspring are shown (n = 55). Zero provisions contained visible dead or desiccated offspring.

| Stage | Number | Percentage |
| --- | --- | --- |
| Viable egg | 20 | 36.4 % |
| Viable larvae | 31 | 56.4 % |
| No offspring | 4 | 7.3 % |

**Table S6:**
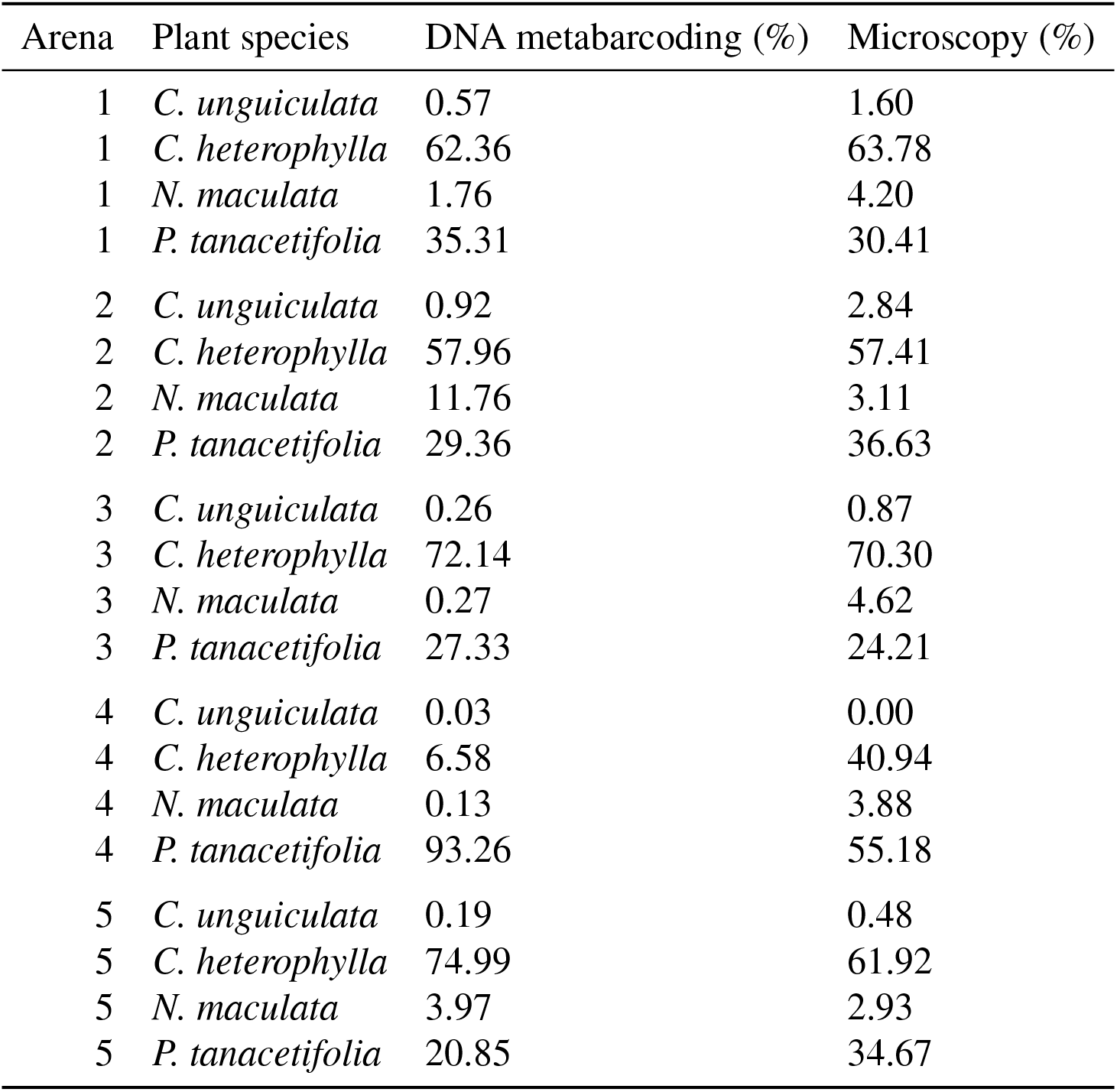
Relative abundance (%) of focal plant species in pollen provisions estimated by ITS2 DNA metabarcoding and microscopy within each experimental foraging arena.

| Arena | Plant species | DNA metabarcoding (%) | Microscopy (%) |
| --- | --- | --- | --- |
| 1 | *C. unguiculata* | 0.57 | 1.60 |
| 1 | *C. heterophylla* | 62.36 | 63.78 |
| 1 | *N. maculata* | 1.76 | 4.20 |
| 1 | *P. tanacetifolia* | 35.31 | 30.41 |
| 2 | *C. unguiculata* | 0.92 | 2.84 |
| 2 | *C. heterophylla* | 57.96 | 57.41 |
| 2 | *N. maculata* | 11.76 | 3.11 |
| 2 | *P. tanacetifolia* | 29.36 | 36.63 |
| 3 | *C. unguiculata* | 0.26 | 0.87 |
| 3 | *C. heterophylla* | 72.14 | 70.30 |
| 3 | *N. maculata* | 0.27 | 4.62 |
| 3 | *P. tanacetifolia* | 27.33 | 24.21 |
| 4 | *C. unguiculata* | 0.03 | 0.00 |
| 4 | *C. heterophylla* | 6.58 | 40.94 |
| 4 | *N. maculata* | 0.13 | 3.88 |
| 4 | *P. tanacetifolia* | 93.26 | 55.18 |
| 5 | *C. unguiculata* | 0.19 | 0.48 |
| 5 | *C. heterophylla* | 74.99 | 61.92 |
| 5 | *N. maculata* | 3.97 | 2.93 |
| 5 | *P. tanacetifolia* | 20.85 | 34.67 |

**Figure S1:**
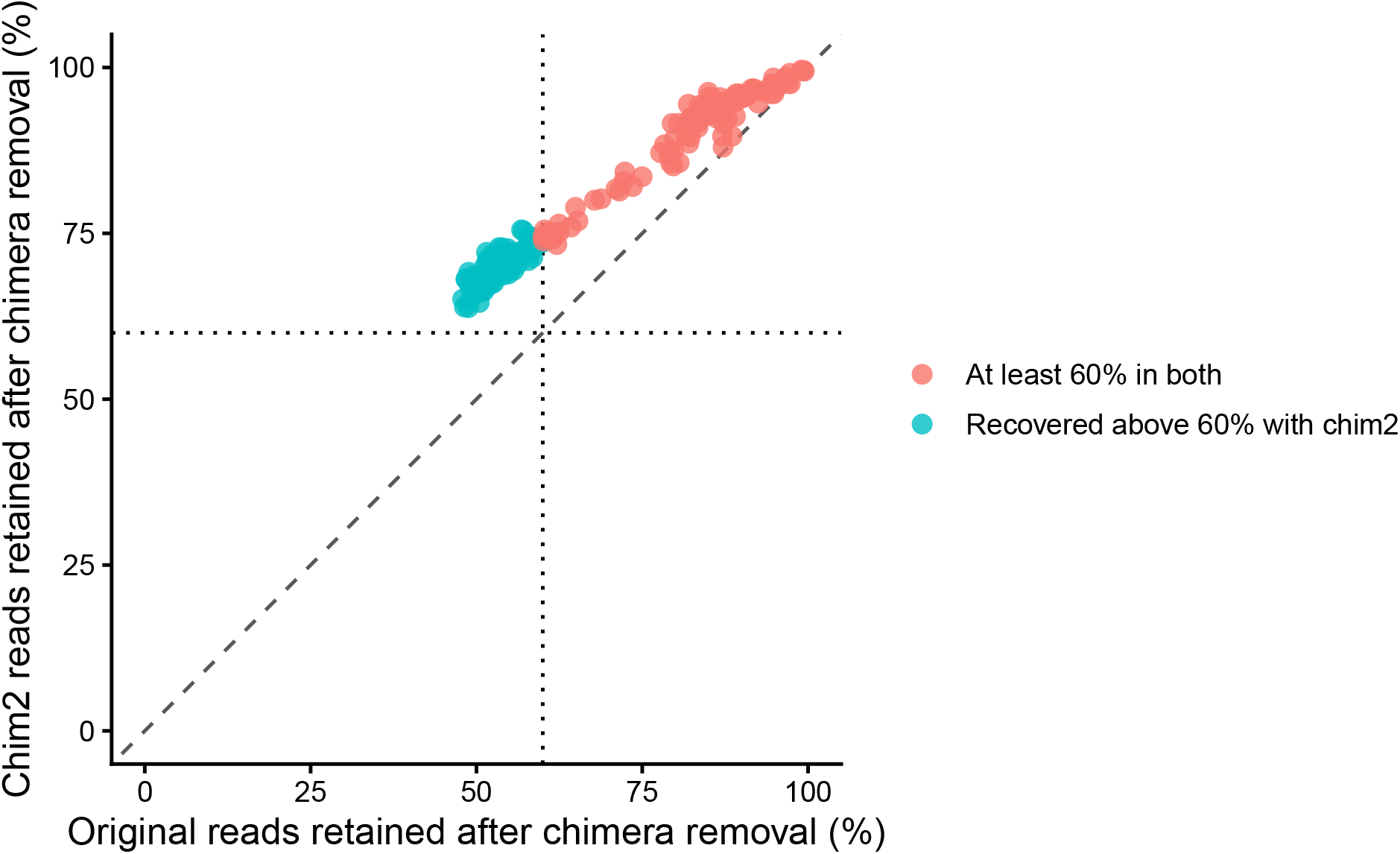
Sample-level sensitivity of read retention to the DADA2 chimera threshold. Each point represents one library. The horizontal axis shows the percentage of merged reads retained after chimera removal using the ratio-one (original) setting minimum parent-over-abundance, and the vertical axis shows retention using a ratio of two. The dashed diagonal denotes equal retention, and dotted lines mark 60% retention. Libraries shown in blue retained less than 60% under the ratio-one (original) setting but more than 60% under the ratio-two setting.

**Figure S2:**
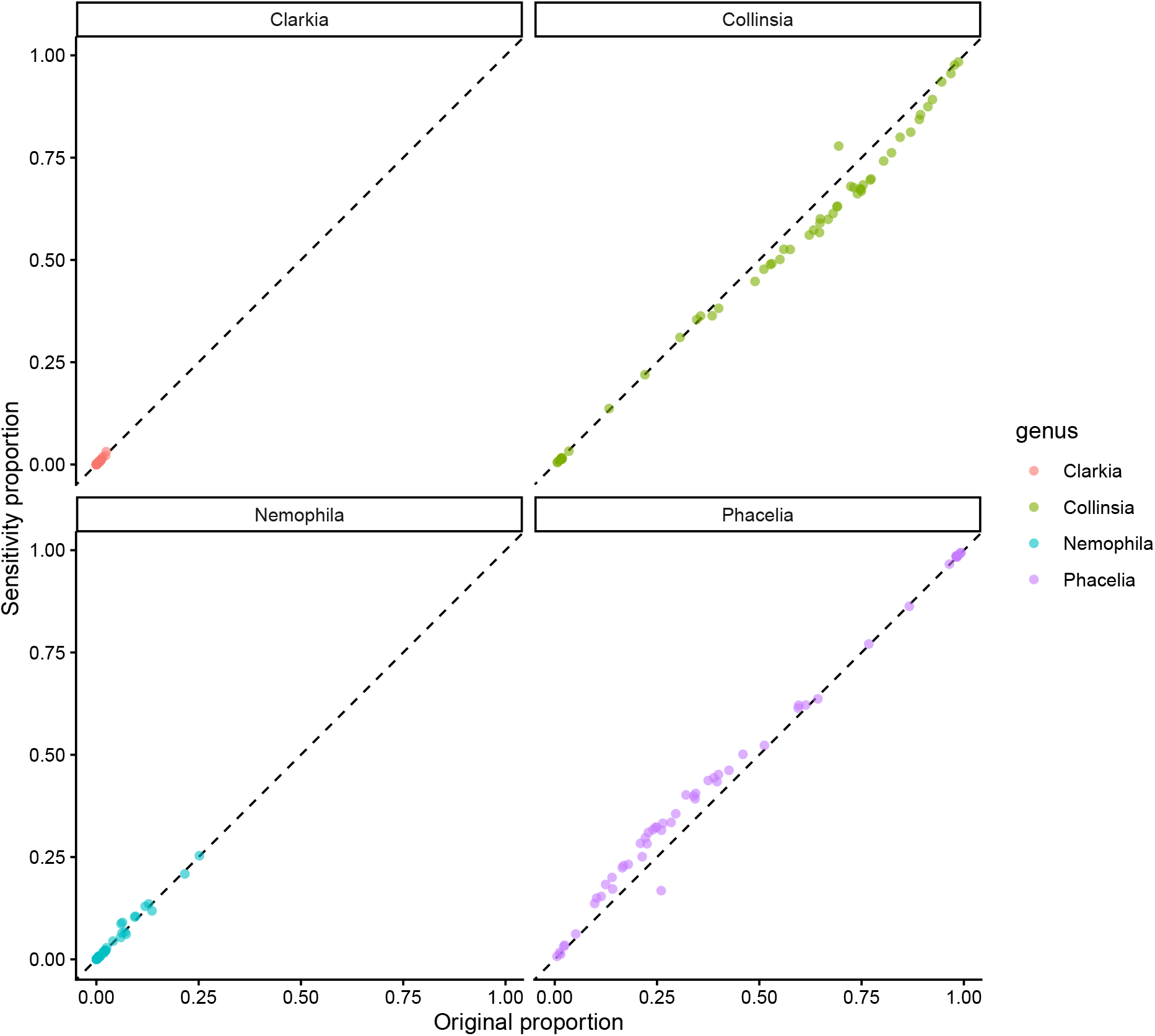
Sensitivity of focal-plant composition to the DADA2 chimera threshold. Relative abundances estimated under the ratio-one setting (original) are plotted against estimates from the ratio-two analysis for matched pollen provisions. Panels show *Clarkia unguiculata*, *Collinsia heterophylla*, *Nemophila maculata*, and *Phacelia tanacetifolia*. The dashed line represents identical relative abundance under both settings. Points clustered near this line show that inferred compositional patterns were robust to the chimera threshold.

**Figure S3:**
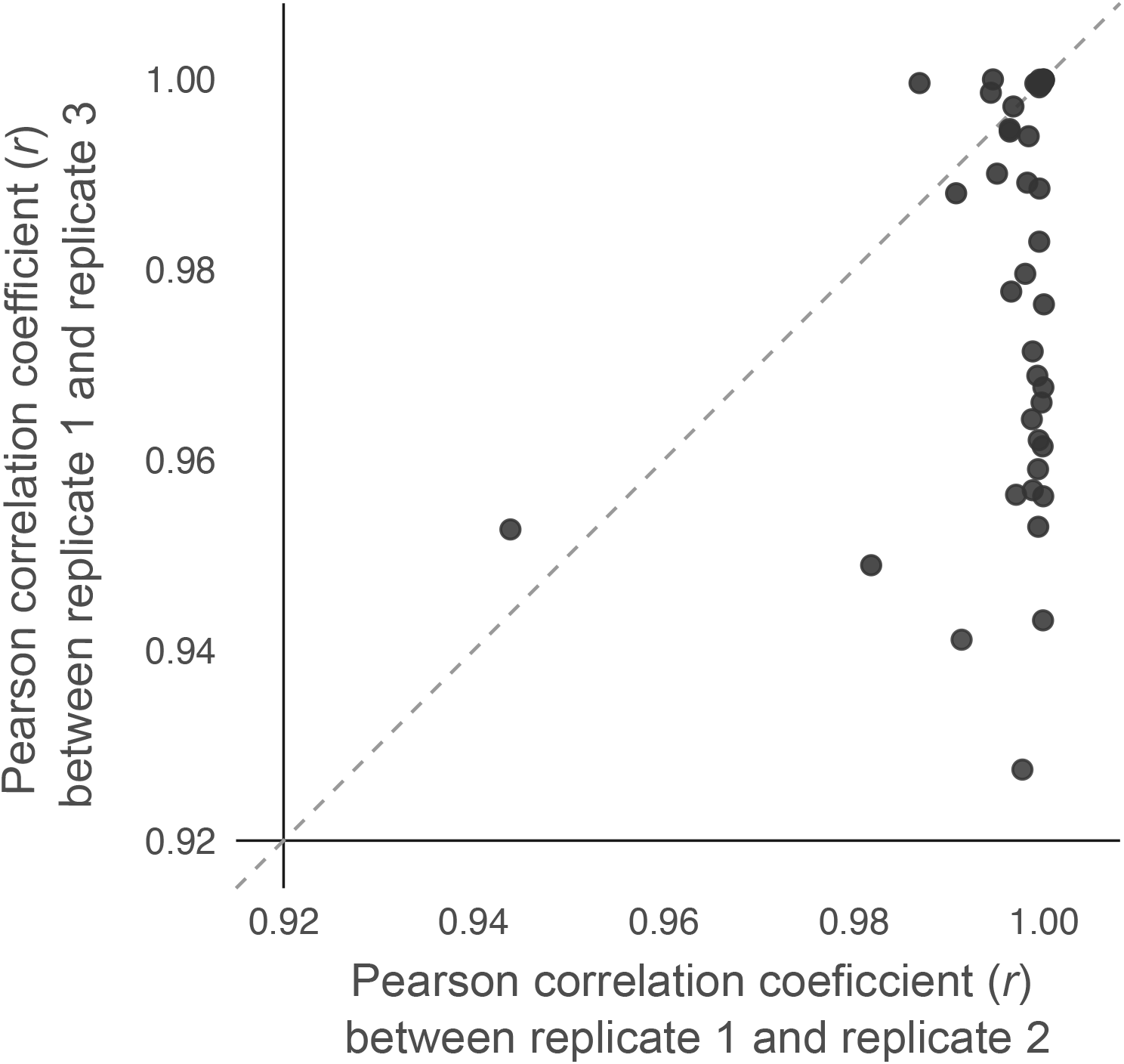
Within-sample consistency using pollen DNA metabarcoding (ITS2). Relationship between each pair of Pearson correlation coefficient for each sample triplicate.

### DADA2 script (qiime2 plugin)

The following script reproduces the DADA2 filtering steps conducted using a minimum fold-parent-over-abundance value of two.

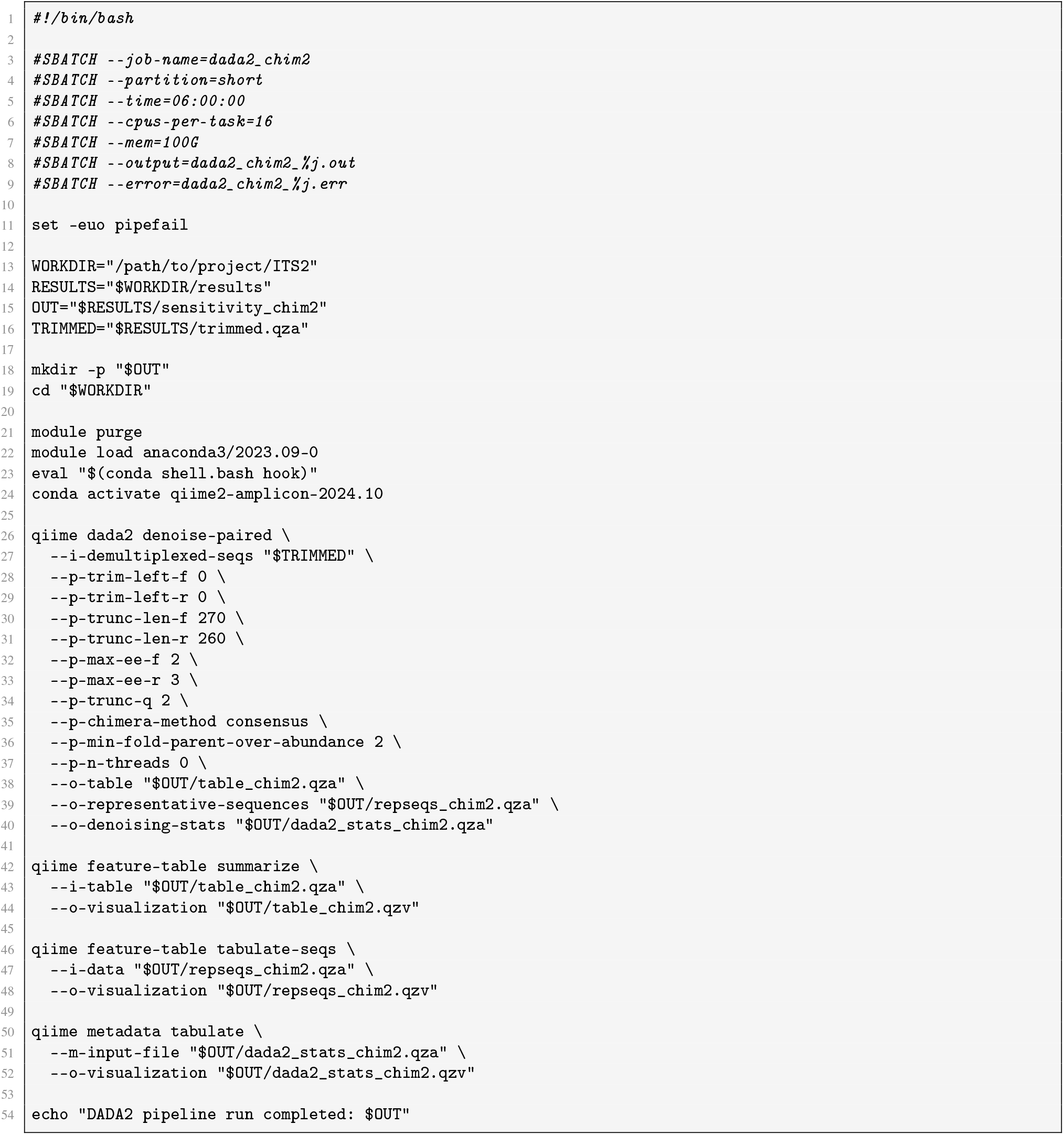

Listing 1: SLURM script for bioinformatics denoising, merging, and chimera detection steps

## Notes

### Competing Interest Statement

The authors have declared no competing interest.

### Summary of Updates

We have revised the focus of our introduction and discussion. We have also added methodological details and improved the supplemental materials.

## References

Altschul, S.F., Gish, W., Miller, W., Myers, E.W. & Lipman, D.J. (1990). Basic local alignment search tool. Journal of Molecular Biology, 215, 403–410.

Anderson, M.J. (2001). A new method for non-parametric multivariate analysis of variance. Austral Ecology, 26, 32–46. _eprint: https://onlinelibrary.wiley.com/doi/pdf/10.1111/j.1442-9993.2001.01070.pp.x.

Andrews, S. (2010). FastQC: A Quality Control tool for High Throughput Sequence Data.

Arstingstall, K.A., DeBano, S.J., Li, X., Wooster, D.E., Rowland, M.M., Burrows, S. & Frost, K. (2021). Capabilities and limitations of using DNA metabarcoding to study plant–pollinator interactions. Molecular Ecology, 30, 5266–5297.

Arstingstall, K.A., DeBano, S.J., Li, X., Wooster, D.E., Rowland, M.M., Burrows, S. & Frost, K. (2023). Investigating the use of pollen DNA metabarcoding to quantify bee foraging and effects of threshold selection. PLOS ONE, 18, e0282715.

Ascher, J.S. & Pickering, J. (2020). Discover Life bee species guide and world checklist (Hymenoptera: Apoidea: Anthophila).

Baksay, S., Andalo, C., Galop, D., Burrus, M., Escaravage, N. & Pornon, A. (2022). Using Metabarcoding to Investigate the Strength of Plant-Pollinator Interactions From Surveys of Visits to DNA Sequences. Frontiers in Ecology and Evolution, 10.

Bell, K.L., Burgess, K.S., Botsch, J.C., Dobbs, E.K., Read, T.D. & Brosi, B.J. (2019). Quantitative and qualitative assessment of pollen DNA metabarcoding using constructed species mixtures. Molecular Ecology, 28, 431–455. _eprint: https://onlinelibrary.wiley.com/doi/pdf/10.1111/mec.14840.

Bolnick, D.I., Svanbäck, R., Fordyce, J.A., Yang, L.H., Davis, J.M., Hulsey, C.D. & Forister, M.L. (2003). The ecology of individuals: incidence and implications of individual specialization. The American Naturalist, 161, 1–28.

Bolyen, E., Rideout, J.R., Dillon, M.R., Bokulich, N.A., Abnet, C.C., Al-Ghalith, G.A., Alexander, H., Alm, E.J., Arumugam, M., Asnicar, F., Bai, Y., Bisanz, J.E., Bittinger, K., Brejnrod, A., Brislawn, C.J., Brown, C.T., Callahan, B.J., Caraballo-Rodríguez, A.M., Chase, J., Cope, E.K., Da Silva, R., Diener, C., Dorrestein, P.C., Douglas, G.M., Durall, D.M., Duvallet, C., Edwardson, C.F., Ernst, M., Estaki, M., Fouquier, J., Gauglitz, J.M., Gibbons, S.M., Gibson, D.L., Gonzalez, A., Gorlick, K., Guo, J., Hillmann, B., Holmes, S., Holste, H., Huttenhower, C., Huttley, G.A., Janssen, S., Jarmusch, A.K., Jiang, L., Kaehler, B.D., Kang, K.B., Keefe, C.R., Keim, P., Kelley, S.T., Knights, D., Koester, I., Kosciolek, T., Kreps, J., Langille, M.G.I., Lee, J., Ley, R., Liu, Y.X., Loftfield, E., Lozupone, C., Maher, M., Marotz, C., Martin, B.D., McDonald, D., McIver, L.J., Melnik, A.V., Metcalf, J.L., Morgan, S.C., Morton, J.T., Naimey, A.T., Navas-Molina, J.A., Nothias, L.F., Orchanian, S.B., Pearson, T., Peoples, S.L., Petras, D., Preuss, M.L., Pruesse, E., Rasmussen, L.B., Rivers, A., Robeson, M.S., Rosenthal, P., Segata, N., Shaffer, M., Shiffer, A., Sinha, R., Song, S.J., Spear, J.R., Swafford, A.D., Thompson, L.R., Torres, P.J., Trinh, P., Tripathi, A., Turnbaugh, P.J., Ul-Hasan, S., van der Hooft, J.J.J., Vargas, F., Vázquez-Baeza, Y., Vogtmann, E., von Hippel, M., Walters, W., Wan, Y., Wang, M., Warren, J., Weber, K.C., Williamson, C.H.D., Willis, A.D., Xu, Z.Z., Zaneveld, J.R., Zhang, Y., Zhu, Q., Knight, R. & Caporaso, J.G. (2019). Reproducible, interactive, scalable and extensible microbiome data science using QIIME 2. Nature Biotechnology, 37, 852–857. Publisher: Nature Publishing Group.

Bosch, J. & Kemp, W.P. (2001). *How to manage the blue orchard bee: as an orchard pollinator*. No. bk. 5 in Sustainable Agriculture Network handbook series. Sustainable Agriculture Network, Beltsville, MD.

Bray, J.R. & Curtis, J.T. (1957). An Ordination of the Upland Forest Communities of Southern Wisconsin. Ecological Monographs, 27, 325–349. _eprint: https://esajournals.onlinelibrary.wiley.com/doi/pdf/10.2307/1942268.

Brooks, M.E., Kristensen, K., van Benthem, K.J., Magnusson, A., Berg, C.W., Nielsen, A., Skaug, H.J., Mächler, M. & Bolker, B.M. (2017). glmmtmb balances speed and flexibility among packages for zero-inflated generalized linear mixed modeling. The R Journal, 9, 378–400.

Burkle, L.A., Marlin, J.C. & Knight, T.M. (2013). Plant-Pollinator Interactions over 120 Years: Loss of Species, Co-Occurrence, and Function. Science, 339, 1611–1615.

Callahan, B.J., McMurdie, P.J., Rosen, M.J., Han, A.W., Johnson, A.J.A. & Holmes, S.P. (2016). DADA2: High-resolution sample inference from Illumina amplicon data. Nature Methods, 13, 581–583. Publisher: Nature Publishing Group.

Camacho, C. (2024). Building a BLAST database with your (local) sequences. In: BLAST® Command Line Applications User Manual [Internet]. National Center for Biotechnology Information (US).

Camacho, C., Coulouris, G., Avagyan, V., Ma, N., Papadopoulos, J., Bealer, K. & Madden, T.L. (2009). BLAST+: architecture and applications. BMC bioinformatics, 10, 421.

Cane, J.H. (2016). Adult pollen diet essential for egg maturation by a solitary Osmia bee. Journal of Insect Physiology, 95, 105–109.

CaraDonna, P.J., Petry, W.K., Brennan, R.M., Cunningham, J.L., Bronstein, J.L., Waser, N.M. & Sanders, N.J. (2017). Interaction rewiring and the rapid turnover of plant–pollinator networks. Ecology Letters, 20, 385–394.

Casanelles-Abella, J., M"uller, S., Keller, A., Aleixo, C., Al’os Orti, M., Chiron, F., Deguines, N., Hallikma, T., Laanisto, L., Pinho, P., Samson, R., Tryjanowski, P., van Mensel, A., Pellissier, L. & Moretti, M. (2022). How wild bees find a way in european cities: Pollen metabarcoding unravels multiple feeding strategies and their effects on distribution patterns in four wild bee species. Journal of Applied Ecology, 59, 457–470.

Chen, S., Yao, H., Han, J., Liu, C., Song, J., Shi, L., Zhu, Y., Ma, X., Gao, T., Pang, X., Luo, K., Li, Y., Li, X., Jia, X., Lin, Y. & Leon, C. (2010). Validation of the ITS2 Region as a Novel DNA Barcode for Identifying Medicinal Plant Species. PLOS ONE, 5, e8613. Publisher: Public Library of Science.

Crone, M.K., Boyle, N.K., Bresnahan, S.T., Biddinger, D.J., Richardson, R.T. & Grozinger, C.M. (2023). More than mesolectic: Characterizing the nutritional niche of Osmia cornifrons. Ecology and Evolution, 13, e10640.

Crone, M.K., Fornoff, F., Klein, A.M. & Grozinger, C.M. (2025). DNA metabarcoding reveals unexpected diet breadth of the specialist large-headed resin bee (Heriades truncorum) in urbanised areas across germany. Insect Conservation and Diversity, 18, 149–160.

Daluwatta Galappaththige, H.S.S. (2024). Approaches to measuring predation pressure. Animal Behaviour, 218, 23–35.

Danforth, B.N., Minckley, R.L., Neff, J.L. & Fawcett, F. (2019). The Solitary Bees: Biology, Evolution, Conservation. Princeton University Press, Princeton, NJ.

Elton, C.S. (1927). Animal ecology. Macmillan Co, New York. Pages: 1–256.

Encinas-Viso, F., Bovill, J., Albrecht, D.E., Florez-Fernandez, J., Lessard, B., Lumbers, J., Rodriguez, J., Schmidt-Lebuhn, A., Zwick, A. & Milla, L. (2023). Pollen DNA metabarcoding reveals cryptic diversity and high spatial turnover in alpine plant–pollinator networks. Molecular Ecology, 32, 6377–6393.

Ewels, P., Magnusson, M., Lundin, S. & Käller, M. (2016). MultiQC: summarize analysis results for multiple tools and samples in a single report. Bioinformatics, 32, 3047–3048.

Fox, G.A., Negrete-Yankelevich, S. & Sosa, V.J. (eds.) (2015). *Ecological Statistics: Contemporary theory and application*. 1st edn. Oxford University PressOxford.

Fründ, J., Linsenmair, K.E. & Blüthgen, N. (2010). Pollinator diversity and specialization in relation to flower diversity. Oikos, 119, 1581–1590.

Gregor, M.I. (2025). Variation in nectar rewards under water limitation across flowering species. Undergraduate Honors Theses, University of San Francisco.

Haider, M., Dorn, S. & Müller, A. (2013). Intra- and interpopulational variation in the ability of a solitary bee species to develop on non-host pollen: implications for host range expansion. Functional Ecology, 27, 255–263. _eprint: https://besjournals.onlinelibrary.wiley.com/doi/pdf/10.1111/1365-2435.12021.

Harano, K.i. & Sasaki, T. (2024). Fuel provisioning for pollen collection by solitary bee, Andrena taraxaci orienticola. Apidologie, 55, 59.

Hartig, F. (2026). DHARMa: Residual Diagnostics for Hierarchical (Multi-Level / Mixed) Regression Models. R package version 0.5.0.

Hengeveld, G., van Langevelde, F., Groen, T. & de Knegt, H. (2009). Optimal Foraging for Multiple Resources in Several Food Species. The American Naturalist, 174, 102–110. Publisher: The University of Chicago Press.

Kaiser-Bunbury, C.N. & Blüthgen, N. (2015). Integrating network ecology with applied conservation: a synthesis and guide to implementation. AoB PLANTS, 7, plv076.

Kembel, S.W., O’Connor, T.K., Arnold, H.K., Hubbell, S.P., Wright, S.J. & Green, J.L. (2014). Relationships between phyllosphere bacterial communities and plant functional traits in a neotropical forest. Proceedings of the National Academy of Sciences, 111, 13715–13720. Publisher: Proceedings of the National Academy of Sciences.

Klečka, J., Mikát, M., Koloušková, P., Hadrava, J. & Straka, J. (2022). Individual-level specialisation and interspecific resource partitioning in bees revealed by pollen DNA metabarcoding. PeerJ, 10, e13671. Publisher: PeerJ Inc.

Labonté, A., Monticelli, L.S., Turpin, M., Felten, E., Laurent, E., Matejicek, A., Biju-Duval, L., Ducourtieux, C., Vieren, E., Deytieux, V., Cordeau, S., Bohan, D.A. & Vanbergen, A.J. (2023). Individual flowering phenology shapes plant–pollinator interactions across ecological scales affecting plant reproduction. Ecology and Evolution, 13, e9707.

Lofgren, L.A., Uehling, J.K., Branco, S., Bruns, T.D., Martin, F. & Kennedy, P.G. (2019). Genome-based estimates of fungal rDNA copy number variation across phylogenetic scales and ecological lifestyles. Molecular Ecology, 28, 721–730.

Martin, M. (2011). Cutadapt removes adapter sequences from high-throughput sequencing reads. EMBnet.journal, 17, 10–12.

Milla, L., Schmidt-Lebuhn, A., Bovill, J. & Encinas-Viso, F. (2022). Monitoring of honey bee floral resources with pollen DNA metabarcoding as a complementary tool to vegetation surveys. Ecological Solutions and Evidence, 3, e12120. _eprint: https://besjournals.onlinelibrary.wiley.com/doi/pdf/10.1002/2688-8319.12120.

Moerman, R., Vanderplanck, M., Fournier, D., Jacquemart, A. & Michez, D. (2017). Pollen nutrients better explain bumblebee colony development than pollen diversity. Insect Conservation and Diversity, 10, 171–179.

Moore, M.A., Scheible, M.K.R., Robertson, J.B. & Meiklejohn, K.A. (2022). Assessing the lysis of diverse pollen from bulk environmental samples for DNA metabarcoding. Metabarcoding and Metagenomics, 6, e89753. Publisher: Pensoft Publishers.

Nemeth, M.B. & Smith-Huerta, N.L. (1997). The effects of pollen clumping on pollen germination and early tube growth in Clarkia unguiculata (Onagraceae). International Journal of Plant Sciences, 158, 292–297.

Oksanen, J., Simpson, G.L., Blanchet, F.G., Kindt, R., Legendre, P., Minchin, P.R., O’Hara, R.B., Solymos, P., Stevens, M.H.H., Szoecs, E., Wagner, H., Barbour, M., Bedward, M., Bolker, B., Borcard, D., Borman, T., Carvalho, G., Chirico, M., Caceres, M.D., Durand, S., Evangelista, H.B.A., FitzJohn, R., Friendly, M., Furneaux, B., Hannigan, G., Hill, M.O., Lahti, L., Martino, C., McGlinn, D., Ouellette, M.H., Cunha, E.R., Smith, T., Stier, A., Braak, C.J.F.T. & Weedon, J. (2025). vegan: Community Ecology Package.

Pedersen, T.L. (2025). patchwork: The Composer of Plots. R package version 1.3.2.9000.

Pornon, A., Andalo, C., Burrus, M. & Escaravage, N. (2017). DNA metabarcoding data unveils invisible pollination networks. Scientific Reports, 7, 16828.

R Core Team (2025). *R: A Language and Environment for Statistical Computing*. R Foundation for Statistical Computing, Vienna, Austria.

Ronca, S., Ford, C.S., Allanguillaume, J., Szabo, C., Kipling, R. & Wilkinson, M.J. (2023). The value of twinned pollinator-pollen metabarcoding: bumblebee pollination service is weakly partitioned within a UK grassland community. Scientific Reports, 13, 18016.

Roulston, T.H., Cane, J.H. & Buchmann, S.L. (2000). What governs protein content of pollen: Pollinator preferences, pollen–pistil interactions, or phylogeny? Ecological Monographs, 70, 617–643.

Sedivy, C., Müller, A. & Dorn, S. (2011). Closely related pollen generalist bees differ in their ability to develop on the same pollen diet: evidence for physiological adaptations to digest pollen. Functional Ecology, 25, 718–725. _eprint: https://besjournals.onlinelibrary.wiley.com/doi/pdf/10.1111/j.1365-2435.2010.01828.x.

Shirazi, S., Meyer, R.S. & Shapiro, B. (2021). Revisiting the effect of pcr replication and sequencing depth on biodiversity metrics in environmental dna metabarcoding. Ecology and evolution, 11, 15766–15779.

Sickel, W., Ankenbrand, M.J., Grimmer, G., Holzschuh, A., Härtel, S., Lanzen, J., Steffan-Dewenter, I. & Keller, A. (2015). Increased efficiency in identifying mixed pollen samples by meta-barcoding with a dual-indexing approach. BMC Ecology, 15, 20.

Sponsler, D., Iverson, A. & Steffan-Dewenter, I. (2023). Pollinator competition and the structure of floral resources. Ecography, 2023, e06651. _eprint: https://nsojournals.onlinelibrary.wiley.com/doi/pdf/10.1111/ecog.06651.

Stephen, K.W. & Rehan, S.M. (2026). Macronutrient composition in pollen affects development and survival in wild bees. Physiological Entomology.

Stephens, D.W. & Krebs, J.R. (1986). *Foraging Theory*. Princeton University Press.

Swenson, S.J. & Gemeinholzer, B. (2021). Testing the effect of pollen exine rupture on metabarcoding with Illumina sequencing. PLOS ONE, 16, e0245611. Publisher: Public Library of Science.

Tornberg, R. & Reif, V. (2006). Assessing the diet of birds of prey: A comparison of prey items found in nests and images. Ornis Fennica, 84.

Tourbez, C., Gómez-Martínez, C., González-Estévez, M.Á. & Lázaro, A. (2024). Pollen analysis reveals the effects of uncovered interactions, pollen-carrying structures, and pollinator sex on the structure of wild bee-plant networks. Insect Science, 31, 971–988. _eprint: https://onlinelibrary.wiley.com/doi/pdf/10.1111/1744-7917.13267.

Valdovinos, F.S., Brosi, B.J., Briggs, H.M., Moisset de Espanés, P., Ramos-Jiliberto, R. & Martinez, N.D. (2016). Niche partitioning due to adaptive foraging reverses effects of nestedness and connectance on pollination network stability. Ecology Letters, 19, 1277–1286. _eprint: https://onlinelibrary.wiley.com/doi/pdf/10.1111/ele.12664.

Vanderplanck, M., Decleves, S., Roger, N., Decroo, C., Caulier, G., Glauser, G., Gerbaux, P., Lognay, G., Richel, A., Escaravage, N. & Michez, D. (2018). Is non-host pollen suitable for generalist bumblebees? Insect Science, 25, 259–272. _eprint: https://onlinelibrary.wiley.com/doi/pdf/10.1111/1744-7917.12410.

Vaudo, A.D., Dyer, L.A. & Leonard, A.S. (2024). Pollen nutrition structures bee and plant community interactions. Proceedings of the National Academy of Sciences, 121, e2317228120.

Vázquez, D.P., Melián, C.J., Williams, N.M., Blüthgen, N., Krasnov, B.R. & Poulin, R. (2007). Species abundance and asymmetric interaction strength in ecological networks. Oikos, 116, 1120–1127.

Weinman, L.R., Ress, T., Gardner, J. & Winfree, R. (2023). Individual bee foragers are less-efficient transporters of pollen for plants from which they collect the most pollen in their scopae. American Journal of Botany, 110, e16178. _eprint: https://bsapubs.onlinelibrary.wiley.com/doi/pdf/10.1002/ajb2.16178.

Westreich, L.R. & Tobin, P.C. (2025). Effect of protein and lipids in pollen on the developmental success of the native solitary bee Osmia lignaria Say. Journal of Apicultural Research, 64, 765–773. _eprint: 10.1080/00218839.2024.2336378.

White, T.J., Bruns, T., Lee, S. & Taylor, J.W. (1990). Amplification and direct sequencing of fungal ribosomal rna genes for phylogenetics. In: PCR Protocols: A Guide to Methods and Applications (eds. Innis, M.A., Gelfand, D.H., Sninsky, J.J. & White, T.J.). Academic Press, New York, pp. 315–322.

Wickham, H., Averick, M., Bryan, J., Chang, W., McGowan, L.D., François, R., Grolemund, G., Hayes, A., Henry, L., Hester, J., Kuhn, M., Pedersen, T.L., Miller, E., Bache, S.M., Müller, K., Ooms, J., Robinson, D., Seidel, D.P., Spinu, V., Takahashi, K., Vaughan, D., Wilke, C., Woo, K. & Yutani, H. (2019). Welcome to the tidyverse. Journal of Open Source Software, 4, 1686.

